# DNA damage and oxidizing conditions activate p53 through differential upstream signaling pathways

**DOI:** 10.1101/2020.09.13.295220

**Authors:** Tao Shi, Paulien E. Polderman, Boudewijn M.T. Burgering, Tobias B. Dansen

## Abstract

Stabilization and activation of the p53 tumour suppressor are triggered in response to various cellular stresses, including DNA damaging agents and elevated Reactive Oxygen Species (ROS) like H_2_O_2_. When cells are exposed to exogenously added H_2_O_2_, ATR/CHK1 and ATM/CHK2 dependent DNA damage signaling is switched on, suggesting that H_2_O_2_ induces both single and double strand breaks. These collective observations have resulted in the widely accepted model that oxidizing conditions lead to DNA damage that subsequently mediates a p53-dependent response like cell cycle arrest and apoptosis. However, H_2_O_2_ induces signaling through stress-activated kinases (SAPK, e.g., JNK and p38MAPK) that can activate p53. Here we dissect to what extent these pathways contribute to functional activation of p53 in response to oxidizing conditions. Collectively, our data suggest that p53 can be activated both by SAPK signaling and the DDR independently of each other, and which of these pathways is activated depends on the type of oxidant used. This implies that it could in principle be possible to modulate redox signaling to stimulate p53 without inducing collateral DNA damage, thereby limiting mutation accumulation in both healthy and tumor tissues.

## 1. Introduction

p53 transcriptional activity induces a wide range of cellular processes including cell cycle arrest, DNA damage repair, senescence, apoptosis and metabolism. Collectively these programs ensure genome integrity, lower the chance to pass on DNA mutations down the lineage and hence provide tumor suppressive function. Nevertheless, p53-dependent programs such as transient cell cycle arrest and the regulation of metabolism can also function to support cell survival, for instance upon nutrient depletion, by providing means to maintain cellular energy levels and control redox balance **[1]**

Under basal, unstressed conditions p53 activity is low as a result of the continuous turnover over p53 protein, which is under control of MDM2 dependent poly-ubiquitinylation and subsequent proteasomal degradation. Stabilization of p53 is the outcome of several cellular stress signaling pathways. Upon DNA double strand breaks, ATM undergoes activating autophosphorylation and phosphorylates p53 on Ser15, but also activates CHK2 which in turn phosphorylates p53 on Ser20 **[2, 3]**. These two phosphorylation events facilitate p53 stabilization by preventing MDM2-mediated p53 ubiquitinylation and subsequent proteasomal degradation **[4]**. In addition to ATM kinase, two members of the stress-activated protein kinase (SAPK) family: c-Jun N-terminal kinase (JNK) and p38MAPK have also been shown to mediate p53 activation in response to UV irradiation, some chemotherapeutic agents but also upon exposure to Reactive Oxygen Species like H_2_O_2_, all of which also have been shown to induce DNA damage **[5–7]**. JNK has been shown to phosphorylate p53 on Ser 20 and Thr 81**[8, 9]**, whereas p38MAPK has been implicated in phosphorylation on Ser15, 33, 37 and 46 **[5, 7]**. Because JNK and p38MAPK are both proline-directed Ser/Thr protein kinases, it may be difficult to pinpoint whether and which of these kinases specifically target a certain site. In any case, these PTMs also induce p53 stabilization and transcriptional activity.

As mentioned, many treatments that engage the cellular DNA damage response also activate SAPK signaling and *vice versa*. It is therefore often difficult to pinpoint which of these pathways is the predominant activator of p53 in many studies **[10, 11]**. For Reactive Oxygen Species (ROS), such as superoxide anions (O_2_^•−^), hydroxyl radicals (HO^•^), hydrogen peroxide (H_2_O_2_), the classical view is that these indeed contribute to damage to proteins, lipids and DNA **[12].** Exogenously added H_2_O_2_ induces both the DNA damage response pathways associated with single and double DNA breaks **[13–15]**. Based on these observations it has been suggested that H_2_O_2_ that is generated endogenously as a consequence of for instance mitochondrial respiration can directly contribute to mutations in genomic DNA, and therefore could be a driver of aging and tumor initiation and progression **[16, 17].**

ROS induced SAPK activation occurs independent of DNA damage as a result of redox signaling: a form of signal transduction that proceeds through the reversible oxidation of protein cysteine-thiols **[18]**. H_2_O_2_ leads for instance to disulfide-dependent dimerization and activation of ASK-1, which activates JNK and p38MAPK which subsequently stabilize p53 **[19, 20].** To complicate things further, ATM has also been reported to be activated by redox signaling independent of DNA DSBs **[21]**. Taken together, and as we recently outlined in detail**[10]**, it remains unclear which upstream signaling pathways (ATM, JNK and p38MAPK) are responsible for oxidant-induced p53 activation in response to which signaling pathways (DNA damage or redox signaling, or both) and to what extent.

In the present study, we aim to dissect signaling cascades upstream of p53 in response to DNA damage signaling and redox signaling. We show that p53 activation in response to DNA damage is mainly mediated by ATM kinase, whereas redox signaling-mediated p53 activation depends on p38MAPK and is independent of the ATM-dependent DNA damage response. ATM, JNK and p38MAPK are all activated by H_2_O_2_, but only ATM and JNK are required for H_2_O_2_-induced p53 activation. The thiol oxidant diamide activates both JNK and p38MAPK but not ATM, and p53 activation by diamide depends on p38MAPK. Collectively, we show that functional p53 activation by redox signaling and DNA damage is mediated by distinct signaling pathways. Our observations imply that for therapeutic strategies p53 can in principle be reactivated by redox signaling without collateral DNA damage, lowering the chance of inducing mutations that drive tumor progression or initiate new malignancies in healthy neighboring tissue.

## 2. Materials and methods

### 2.1 Reagents and antibodies

Diamide, H_2_O_2_, Neocarzinostatin (NCS), Auranofin (AFN), and ATM inhibitor (KU55933) were from Sigma. JNK inhibitor (SP600126) and p38MAPK inhibitor (PH797804) were from Bio-Connect Life Sciences. Nutlin-3a was from Sanbio.

Antibodies were used as follows: p53(DO-1), p21(M-19), JNK (D-2), c-Jun and p38MAPK were from Santa Cruz Biotechnology. Anti-pp53 Ser15, Ser20, pCHK2(Thr68), p-C-Jun (Ser63), pJNK (Thr183/Tyr185), pATM (Ser1981), pp38MAPK (Thr180/Tyr182) and pATF-2 (Thr71) were from Cell Signaling Technology. Phospho-Histone H2AX (Ser139) and GAPDH (MAB374) were from EMD Millipore. HRP or fluorescently labeled secondary antibodies were used for detection on Western blot.

### 2.2 Cell culture

HEK293T cells and non-small-cell-lung cancer cells (NCI-H1299, ATCC^®^ CRL-5803™) **[22]** cells were cultured in DMEM high-glucose (4,5g/L) supplemented with 10 % FBS, 2mM L-glutamine and 100 Units Penicillin-Streptomycin (All from Sigma Aldrich). RPE^Tert^ cells were cultured in DMEM/F-12 high-glucose supplemented with 10 % FBS and 100 U Penicillin-Streptomycin (Sigma Aldrich). All cell types were cultured at 37 °C under a 6 % CO_2_ atmosphere. Cell transfection was carried out using PEI (Sigma Aldrich).

p53-KO RPE^Tert^ cells were a gift from R.Medema **[23]**. Stable, Doxycycline-inducible p53 expressing cells were generated by transduction with lentiviral pInducer20-Flag-p53 in the p53-KO RPE^Tert^ or H1299 background, followed by selection with 400 μg/ml (for RPE^Tert^ cells) and 600 μg/ml (for H1299 cells) Neomycin for 2 weeks. pInducer20 plasmid was a gift from Stephen Elledge (Addgene plasmid # 44012; http://n2t.net/addgene:44012; RRID: Addgene_44012). pInducer20-Flag-p53 was made through Gateway cloning following standard procedures **[24]**. The inducible expression of p53 was confirmed by Western blotting and polyclonal cells were used for subsequent experiments.

### 2.3 Western Blotting

RPE^Tert^ or H1299 cells were seeded in 6-well dishes and growing to be around 80 % confluency, followed by treatments with different compounds for the indicated time. Cells were then directly scraped in 1X sample buffer (2 % SDS, 5 % 2-mercaptoethanol, 10 % glycerol, Tris-HCI pH 6.8, 0.002 % bromophenol blue). Samples were run on SDS-PAGE gels (Biorad system), followed by a standard Western blotting precure. Briefly, samples were transferred to polyvinylidene difluoride (PVDF) or nitrocellulose membranes. Membranes were then blocked with 2 % BSA TBS-Tween (TBST, 1 % v/v) solution for 1h at 4 °C, followed by incubation with primary antibodies overnight at 4 °C. After washing the membrane with TBST solution, secondary antibody staining was performed using HRP or fluorescence-conjugated antibodies for 1h at 4 °C. After washed three times with TBST, membranes were analyzed by Image Quant LAS or Typhoon-Biomolecular Imager.

### 2.4 Ubiquitinylation assay

HEK293T cells were transiently transfected with Flag-His-p53 and His-ubiquitin expression constructs. After 48 h, cells were treated with H_2_O_2_ or diamide (15 min) followed by lysis in buffer containing 100 mM NaH_2_PO_4_/Na_2_HPO_4_, 10 mM Tris pH 8, 8 M Urea, 10 mM Imidazole and 0.2 % Triton X-100 and sonication. Cell lysates were then centrifuged at 10,000 rpm for 10 min, and 50 μl of supernatant was taken as input sample and ubiquitinated proteins were enriched by incubation with Ni-NTA beads for 2 h at room temperature. The Ni-NTA beads were washed twice with the above indicated lysis buffer, followed by a wash with elution buffer (100 mM NaCl, 20 % glycerol, 20 mM Tris pH 8.0, 1mM DTT and 10 mM Imidazole). In the end, ubiquitinated proteins were resuspended in 1X sample buffer (2 % SDS, 5 % 2-mercaptoethanol, 10 % glycerol, Tris-HCI pH 6.8, 0.002 % bromophenol blue), boiled at 95 °C for 8 mins, and further analyzed by standard Western blotting.

### 2.5 Immunofluorescence microscopy

RPE^Tert^ cells were grown on glass coverslips in 6-well dishes for three days and then treated with PBS, diamide, H_2_O_2_ or NCS for 1 h. Cells were washed twice with cold PBS and fixed (3.7 % Formaldehyde solution) for 15 min at room temperature. Fixed cells were permeabilized using 0.1 % Triton for 5 min followed by blocking with 2 % BSA (w/v) plus purified goat IgG in 1:10,000 in PBS for 30-60 min at room temperature. After that, cells were incubated with primary antibodies (1:500 dilution for Anti-p53 and pH2AX (Ser139)) overnight, followed by 1 h incubation with fluorescently labeled secondary antibodies (1:500) and Hoechst (1:10,000) post washing twice with PBS. All antibody incubations were performed at 4°C and in the dark. Coverslips were mounted with a drop of mounting medium, sealed with nail polish to prevent drying, and saved in the dark at 4 °C until analysis. Imaging was performed on a Zeiss Axio Imager Z1and images were processed using ImageJ software.

### 2.6 RNA isolation and qPCR

Total RNA was isolated from Doxycycline-inducible p53 expressing RPE^Tert^ cells (treated with or without doxycycline) using the RNeasy kit (QIAGEN). 500 ng RNA was used for cDNA synthesis using the iScript cDNA Synthesis Kit (BIO-RAD) according to the manufacturer’s instructions. qPCR experiments were carried out using SYBR Green FastStart Master Mix **[25]** on a CFX Connect Real-time PCR detection system (Bio-RAD). qPCR cycle settings were as follows: pre-denaturation at 95 °C for 10 min, followed by 39 cycles of denaturation at 95 °C for 10 s, annealing at 58 °C for 10 s, and extension at 72 °C for 30 s. Relative gene expression was calculated using 2^−ΔΔCT^ method by taking GAPDH as a reference gene. All the primers used for qPCRs are shown in **Table S1**.

### 2.7 Flow cytometry

To assess the effect of the various treatments on the DNA damage response and the nuclear redox state (measured as the ratio of oxidation and reduction of the H_2_O_2_-specific sensor HyPer2), HEK293T cells were transfected with an H2B-HyPer fusion construct and 48 h later treated with diamide, H_2_O_2_ or NCS for the indicated time (15 min, 30 min, 1 h, 2 h and 6 h). To prevent post-harvest HyPer oxidation or reduction, cells were washed with PBS containing 100 mM NEM prior to trypsinization. Cells were fixed with 4 % Formaldehyde solution for 10 min at room temperature, followed by the incubation with ice-cold 70 % Ethanol overnight at 4 °C. Cells were resuspended in PBS and washed with PBS buffer twice, and stained with anti-Phospho-Histone H2AX (Ser139) PE-conjugated antibody (0.25 ug/sample) for 30 min at room temperature. After that, cells were resuspended in PBS buffer with 1 % BSA and 0.05 % Tween and further analyzed by a BD FACSCelesta Flow Cytometer (BD Bioscience). H2B-HyPer2 fluorescence was detected using Ex405/Em525/50 (reduced) and Ex488/Em30/30 (oxidized) lasers and filtersets.

Cell death was assessed using Propidium Iodide staining. RPE^Tert^ cells and H1299 cells with or without doxycycline were treated with diamide, H_2_O_2_ or NCS for 24 h. Cell medium, PBS buffer used to wash cells and trypsinized cells were collected together. Samples were washed once with PBS and resuspended in PBS containing propidium iodide (PI) (sigma-Aldrich) and incubated for 15-30 min in the dark. Cells were then analyzed on a BD FACSCelesta Flow Cytometer (BD Bioscience). Cells positive for PI were considered dead.

### 2.8 Statistical analysis

All statistical analysis was performed in GraphPad Prism 8 software. One-way ANOVA method followed by Dunnett’s multiple comparisons, was used to evaluate the statistical significance of qPCR data, and an adjusted p value < 0.05 was considered to be statistically significant. A Student’s t-test was used to evaluate the difference of cell death in p53-off and p53-on cells upon each treatment, and a p-value < 0.05 was considered to be statistically significant.

## 3. Results

### 3.1 Differential activation of redox signaling and the DNA damage response

As mentioned in the introduction of this study, H_2_O_2_ is known to induce both redox signaling and the DDR. Likewise, several genotoxic agents used in DNA damage research have been suggested to act, at least in part, through the production of reactive oxygen species and hence could start or modulate redox signaling through a decrease in reductive power. To be able to dissect the DDR and redox signaling-based responses, we investigated whether it is possible to activate these pathways independently. To this end the ratio of oxidized (λ_Ex_488 nm / λ_Em_ 530/30nm) over reduced (λ_Ex_405 nm / λ_Em_ 525/50 nm) HyPer2 **[26]** as well as positivity for H2AX-pSer139 (aka γH2AX) was assessed in HEK293T cells by flow cytometry upon treatment with diamide, H_2_O_2_ or Neocarzinostatin (NCS). The HyPer2 probe was localized to the chromatin by means of a Histone-H2B fusion in order to assess H_2_O_2_/redox signaling in the vicinity of the DNA, which we assumed most relevant for potential oxidation induced DNA damage. **Figure 1** shows that it is indeed possible to induce DDR and oxidizing conditions separately for prolonged time periods. The thiol-specific oxidant diamide, which is thought to act largely through oxidation of the GSH pool, rapidly but transiently induced HyPer2 oxidation, without affecting γH2AX levels for up to 6 hrs after treatment. Note that the ratio of the HyPer2 probe is a measure of the combined rate of oxidation (by H_2_O_2_) and reduction (by the thioredoxin system) **[27]**, which makes it difficult to distinguish whether the diamide-induced increase in HyPer2 ratio stems from an increase of H_2_O_2_ from endogenous sources or from a loss of reductive power or both. In any case, this also means that the ratio of the H_2_O_2_-specific HyPer2 probe correlates with that of PRDX oxidation/reduction and hence is a good read-out for the induction of redox signaling. The DNA damaging agent NCS induced a buildup of γH2AX signal that peaked 1 h after treatment, without evidence of changes in the HyPer2 ratio. Treatment with H_2_O_2_ resulted in both HyPer2 oxidation and phosphorylation of H2AX, in line with the idea that this compound indeed induces both redox signaling and the DDR **[28]**. Note that the kinetics of HyPer2 oxidation and H2AX phosphorylation by H_2_O_2_ follow those of diamide and NCS respectively. The induction of DNA damage by H_2_O_2_ and NCS but not diamide was further corroborated using immunofluorescence microscopy on non-transformed, human Telomerase immortalized Retinal Pigment Epithelial (RPE^Tert^) cells (**Fig. 1D**). Taken together, diamide and NCS can serve as model compounds in this study to dissect to what extent the effects of H_2_O_2_ treatment are mediated through redox signaling, the DDR or both. Furthermore, these data already indicate that oxidizing conditions do not necessarily lead to a DDR.

**Figure 1.**
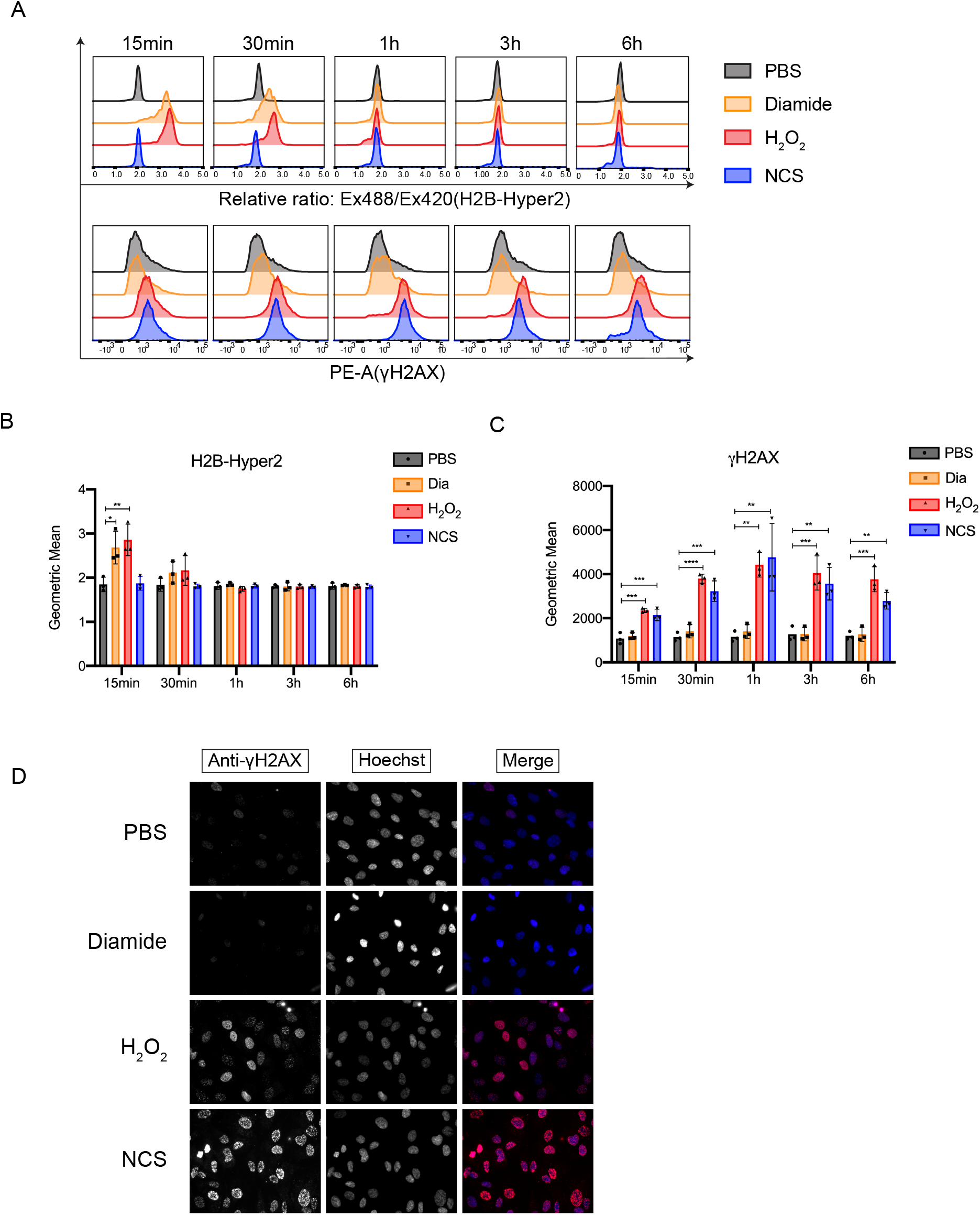
Redox signaling and DNA damage signaling can be induced independent of each other. (A) HEK293T cells transiently expressing H2B-HyPer2 were treated with PBS, diamide (200 μM), H_2_O_2_ (200 μM) or Neocarzinostatin (NCS) (250 ng/ml) for the indicated times. The ratio of oxidized (Ex 488 nm / Em 530/30 nm) over reduced (Ex405 nm / Em 525/50 nm) HyPer2 and the level of γH2AX (H2AX-pSer139) were evaluated by Flow Cytometry. (B) Quantification of geometric mean of the H2B-HyPer2 ratio from three independent experiments. The error bars stand for the standard division (SD) of the geometric means. The H2B-HyPer2 ratio in treated groups (diamide, H_2_O_2_ and NCS) was compared with PBS (CTRL) group. The statistical analysis was performed by using One-way ANOVA followed by Multiple comparisons test (Dunnett) under each time point. *, p value < 0.05; **, p value < 0.005. (C) Quantification of geometric mean of γH2AX from three independent experiments. The error bars stand for the standard division (SD) of the geometric means. The γH2AX intensity in treated groups (diamide, H_2_O_2_ and NCS) was compared with PBS (Ctrl) group. The statistical analysis was performed by using One-way ANOVA followed by Multiple comparisons test (Dunnett) under each time point. *, p value < 0.05; **, p value < 0.005; ***, p value < 0.0005; ****, p value < 0.0001. (D) Epifluorescence microscopy staining for γH2AX in RPE^Tert^ cells upon 1h treatment of diamide (200 μM), H_2_O_2_ (200 μM) or NCS (250 ng/ml). Nuclei were stained with Hoechst and γH2AX were stained with an anti-γH2AX primary antibody and Alexa Fluor 568 as the secondary antibody.

### 3.2 Redox signaling activates p53 independent of the DDR

Now that we had found means to selectively induce oxidizing conditions without triggering the DDR, we set out to explore whether and how this contributes to p53 stabilization and activation upon exposure to H_2_O_2_. To this end, we continued with non-transformed Retinal Pigment Epithelial (RPE^Tert^) cells, which have a more wildtype p53 response. These RPE^Tert^ cells were exposed to diamide or H_2_O_2_ for various timepoints **(Fig. 2A)**. NCS was used as a positive control for DDR activation in the absence of redox signaling (as shown in Fig. 1A). Like observed in HEK293T, treatment with diamide also did not trigger the DDR in RPE^Tert^ cells, as evidenced by the absence of ATM-pS1981, CHK2-pThr68 and p53-pSer15 induction. Nevertheless, prolonged (6 h) treatment with diamide surmounted in p53 stabilization to comparable levels as that induced by NCS (1 h) or H_2_O_2_ (6 h), and this was accompanied by accumulation of the p53 transcriptional target gene product p21. Indeed, p21 was not induced in CRISPR/CAS9-derived RPE^Tert^ p53 KO cells **(Fig. 2B)**. Both oxidizing compounds (but not NCS) trigger JNK (T183/Y185) phosphorylation, albeit more pronounced by diamide, indicating that the SAPK pathway acts downstream of redox signaling and independent of the DDR. Note that the stabilization of p53 by H_2_O_2_ and diamide was observed long after JNK or CHK2 phosphorylation had ceased.

**Figure 2.**
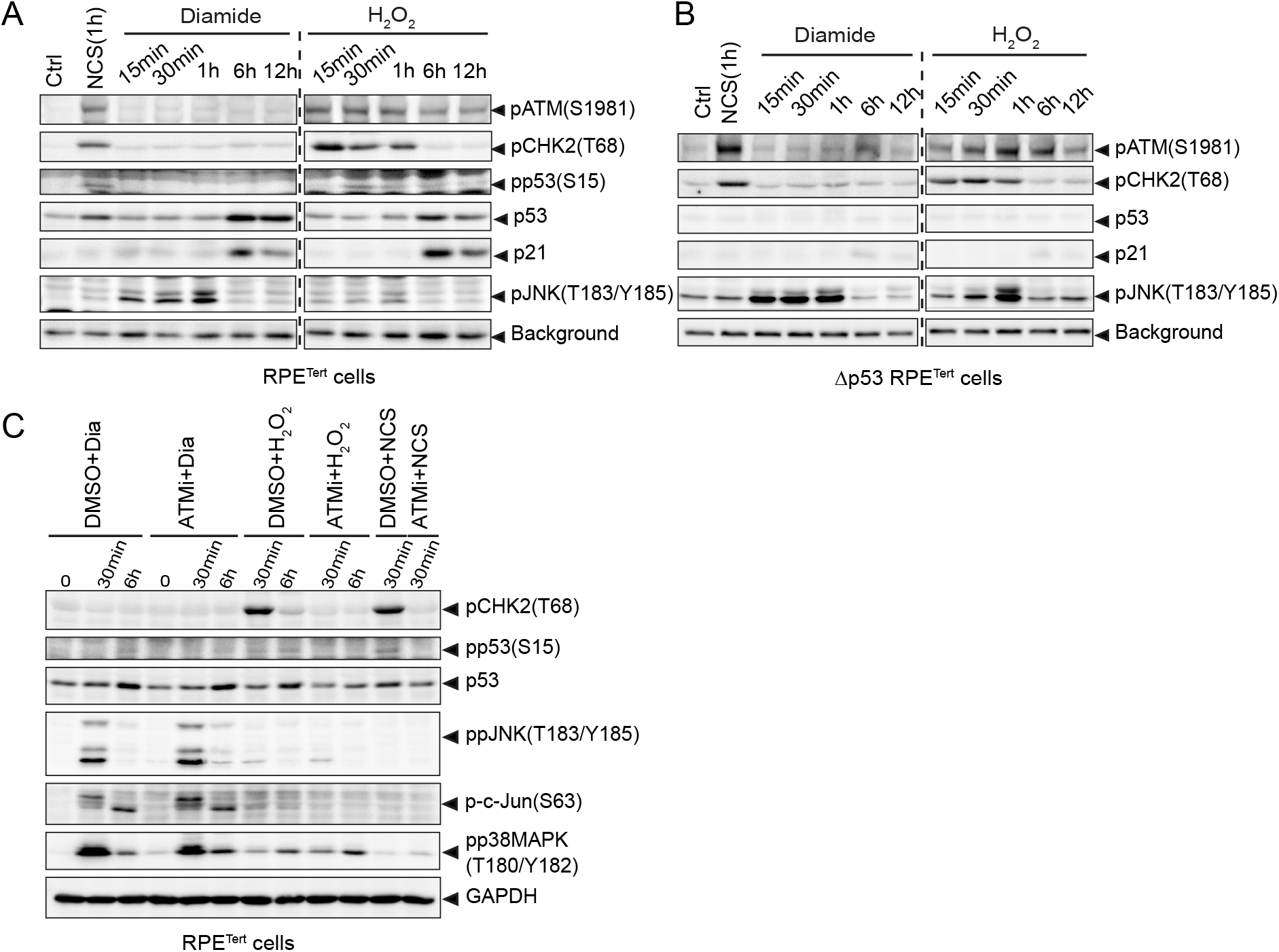
Redox signaling activates p53 independent of the ATM-dependent DNA damage response. (A) RPE^Tert^ cells were treated with NCS (250 ng/ml), diamide (200 μM) and H_2_O_2_ (200 μM) for the indicated time. Endogenous p53 and p21 levels and phosphorylation states of ATM (S1981), CHK2(T68), p53(S15) and JNK (T183/Y185) were evaluated by immunoblotting. (B) Same treatment as in (A), but using RPE^Tert^ p53 KO cells. (C) RPE^Tert^ cells were pretreated with DMSO or ATM inhibitor (ATMi) KU55933 (10 μM) for 1h, followed by treatment with diamide (200 μM), H_2_O_2_ (200 μM) or NCS (250 ng/ml) for the indicated time. p53 level, phosphorylation state of CHK2(T68), p53(S15), JNK(T183/Y185), p38MAPK(T180/Y182) and GAPDH as a loading control were evaluated by immunoblotting.

To further elucidate signaling downstream of the redox signaling and DDR response, we assessed whether ATM activity was required for the observed stabilization of p53 by oxidizing (diamide/H_2_O_2_) versus DNA damaging (H_2_O_2_/NCS) conditions using the ATM inhibitor KU55933 (ATMi). Inhibition of ATM abolished p53 phosphorylation on Ser15 and stabilization induced by H_2_O_2_ and NCS, whereas this had no effect on diamide-induced p53 stabilization (**Fig. 2C**). This suggests that whereas the DDR downstream of H_2_O_2_ proceeds through ATM, redox signaling does not. Phosphorylation of CHK2 was indeed largely abolished by ATMi, whereas both JNK and p38MAPK were unaffected. Like observed in Fig2A, B diamide induced JNK and p38MAPK activation to a larger extent as compared to H_2_O_2_. If redox signaling-induced p53 stabilization depends on SAPK activation, this observation could be an explanation as to why redox signaling downstream of H_2_O_2_ fails to stabilize p53 in the presence of ATMi; something we will explore later in this study. Collectively, these results indicate that the redox signaling and the ATM-dependent DNA damage signaling responses as observed upon H_2_O_2_ exposure can be induced independent of each other, and that both pathways can lead to p53 stabilization and activation.

### 3.3 Diamide and H_2_O_2_ stabilize p53 through inhibition of its ubiquitin-dependent degradation

Several cellular stresses, including DNA damage, have been shown to induce stabilization of p53 through interference with MDM2-dependent ubiquitinoylation and subsequent proteasomal degradation **[29, 30]**. To explore whether diamide and H_2_O_2_-induced p53 stabilization also depend on inhibition of protein breakdown, p53 protein decay dynamics were assessed in the presence of these compounds in combination with the protein synthesis inhibitor cycloheximide (CHX). p53 levels rapidly declined under control conditions and persisted upon treatment with the positive control Nutlin-3a (an MDM2 inhibitor). Treatment with diamide, and to a lesser extent H_2_O_2_, resulted in attenuated p53 decay, suggesting that these oxidants interfere with MDM2-dependent degradation (**Fig. 3A and 3B).** In accordance, ubiquitinylation of p53 was inhibited upon diamide and H_2_O_2_ treatment (**Fig. 3C**). Several enzymes involved in the (de)ubiquitinylation reaction depend on catalytic cysteines and thus may be negatively regulated through oxidation. However, total protein ubiquitinylation appeared unaffected suggesting a specific effect of oxidants on ubiquitin-dependent p53 degradation.

**Figure 3.**
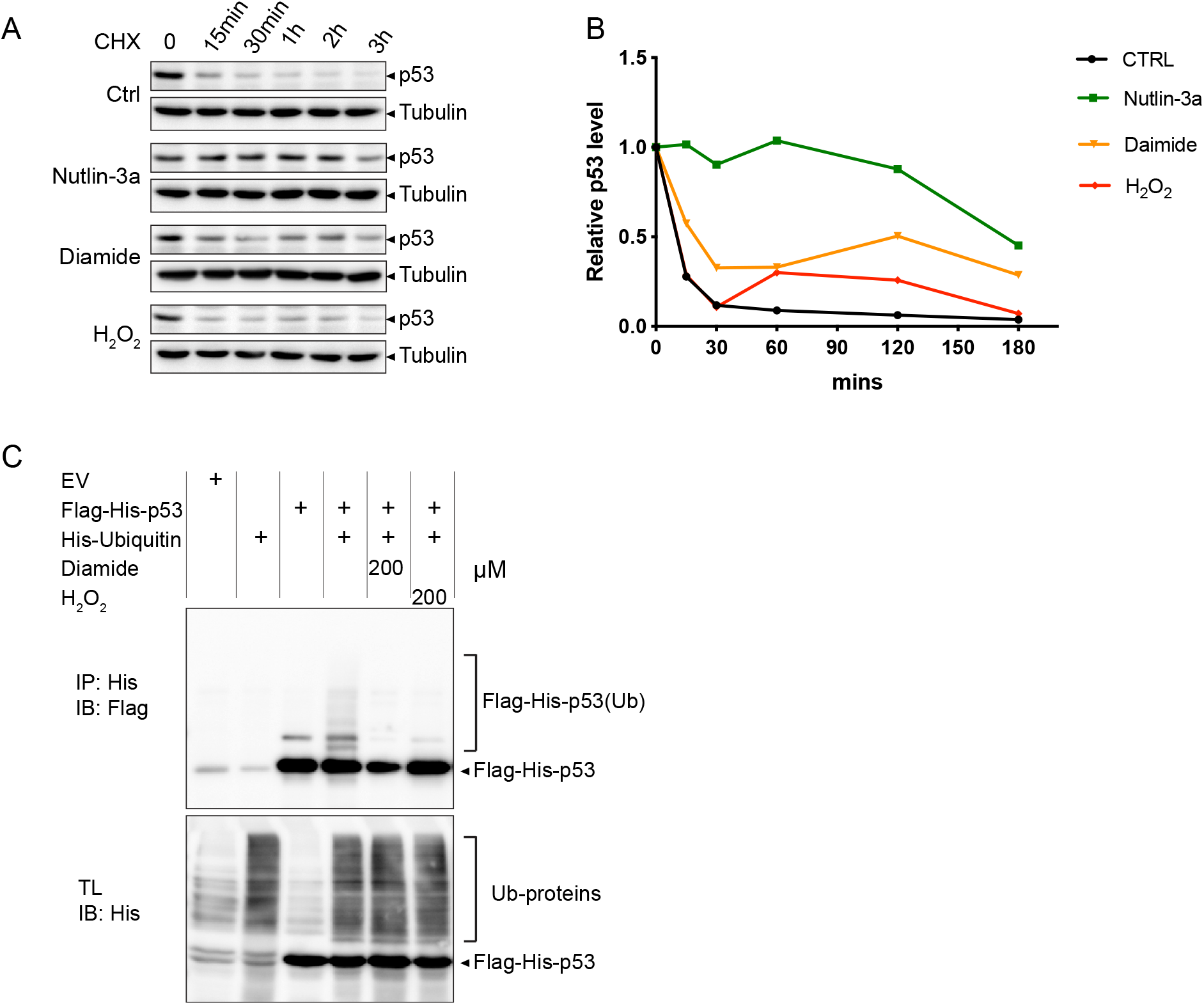
Diamide and H_2_O_2_ stabilize p53 by inhibition of protein degradation and ubiquitinylation. (A) RPE^Tert^ cells were treated with Cycloheximide (CHX, 10 μg/ml) to block protein synthesis and at the same time exposed to Nutlin-3a (10 μM), diamide (200 μM) or H_2_O_2_ (200 μM) for the indicated time. Total cell lysates were loaded for evaluating the levels of endogenous p53 and Tubulin (as a loading control). (B) Quantification of p53 protein intensity relative to Tubulin from (A). (C) HEK293T cells expressing Flag-His-p53 alone or in combination with His-ubiquitin were treated with diamide (200 μM) or H_2_O_2_ (200 μM) for 15 min. p53 ubiquitinylation was evaluated by His-pulldown using Ni-NTA beads and immunoblotting with anti-Flag antibody. The presented data are representative for three independent experiments.

### 3.4 Diamide and H_2_O_2_ dependent p53 activation are mediated by different Stress Activated Protein Kinases (SAPKs)

To further investigate how p53 was stabilized and activated by redox-dependent signaling, we made use of inhibitors of JNK and p38MAPK kinases **(Fig. 4A)**: two SAPKs that have previously been shown to be activated by redox signaling and that have both been implicated in p53 activation **[5, 6, 31]**, Also in our experiments these pathways were activated by both diamide and H_2_O_2_, although diamide generally resulted in a slightly stronger activation (see also earlier in **Fig. 2**). Pre-treatment with the JNK inhibitor SP600125 strongly inhibited JNK activity induced by both diamide and H_2_O_2_ (most notably observed by assessment of its downstream target c-jun-pS63) (**Fig. 4B**). Strikingly, whereas SP600125 pre-treatment almost completely abolished H_2_O_2_ induced p53 stabilization and activation, evidenced by p21 induction, diamide-dependent signaling towards p53 remained unaffected. Conversely, inhibition of p38MAPK by pre-treatment with PH797804 largely blocked diamide-induced p53 stabilization and p21 induction, whereas p53 activation by H_2_O_2_ was not affected by p38MAPK inhibition. Note that the effect of PH797804 on p53 was already noticeable, despite some p38MAPK activity remained as judged by some remaining ATF2-pT71 (a target of p38MAPK) signal (**Fig. 4C**). Pre-treatment with both inhibitors indeed blocked the induction of p53 and p21 induced by either redox signaling stimulus. These observations strongly indicate that even though diamide and H_2_O_2_ both activate p38MAPK and JNK, diamide-dependent p53 activation is mediated by p38MAPK, whereas H_2_O_2_ mediated p53 activation is mediated by JNK. We showed earlier (Fig. 2) that H_2_O_2_ also requires ATM signaling, whereas p38MAPK does not. The experiments using the JNK inhibitor suggest that under these conditions, ATM signaling is still active (induction of CHK2-pT68, **Fig. 4B**), but not sufficient for p53 activation. Apparently both JNK and ATM activity are needed for full activation of p53 by H_2_O_2_ treatment.

**Figure 4.**
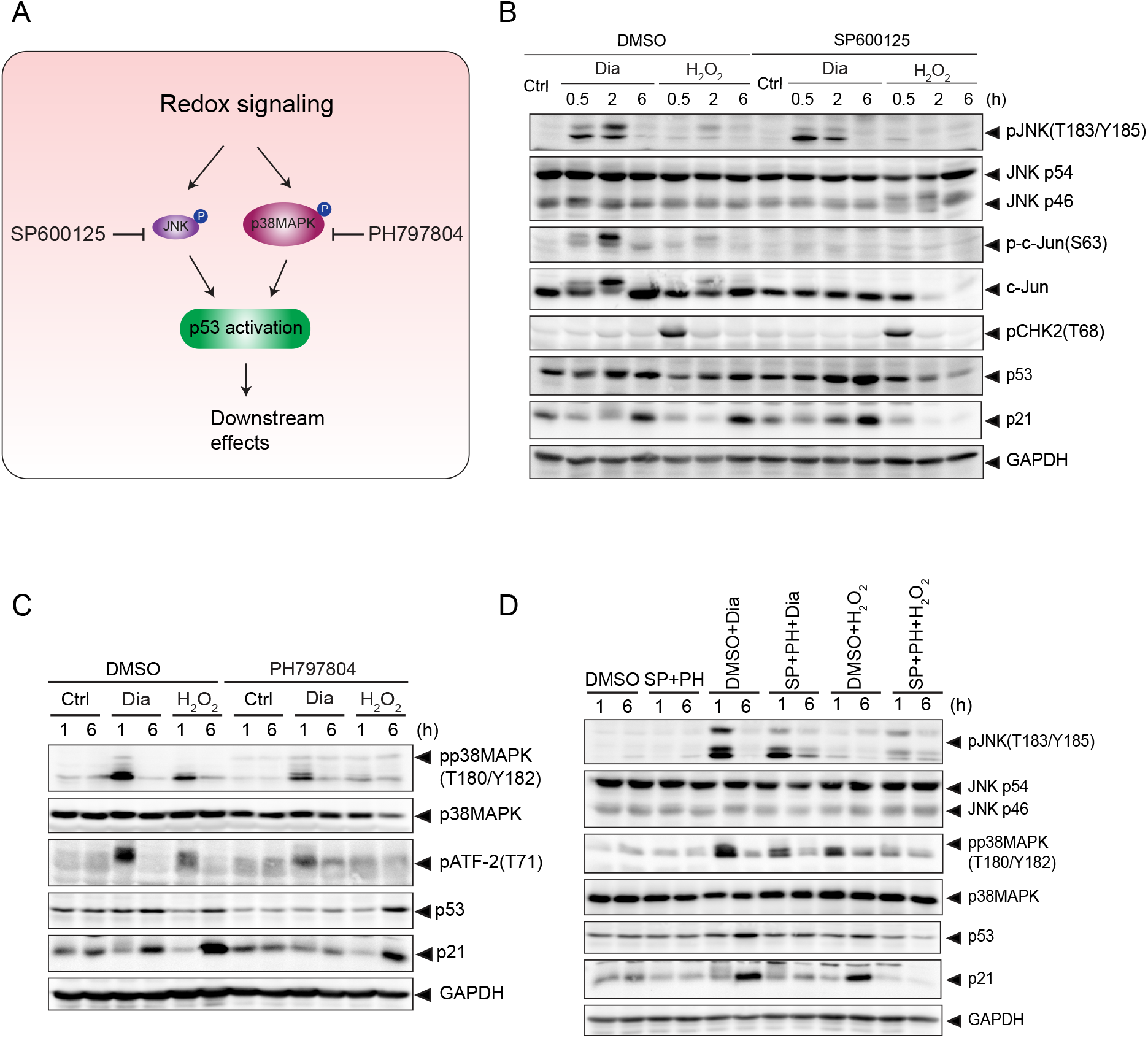
p38MAPK, not JNK, is required for redox signaling-mediated p53 activation. (A) Overview of p53 activation through p38MAPK and JNK under redox signaling. p38MAPK and JNK are activated in response to redox signaling, which leads to p53 activation. SP600125 and PH797804 are inhibitors for JNK and p38MAPK, respectively. (B) JNK is dispensable for diamide-mediated p53 activation, but is essential for H_2_O_2_-induced p53 activation. RPE^Tert^ cells were pre-treated with DMSO or SP600125 (20μM) for 2h, followed by treatment with diamide (200 μM) or H_2_O_2_ (200 μM) for the indicated time. The total protein levels of JNK (p54 and p46), c-Jun, p53, p21 and GAPDH, and phosphorylation states of JNK(T183/Y185), c-Jun(S63) and CHK2(T68) were detected by immunoblotting. (C) p38MAPK is indispensable for diamide-induced, but not for H_2_O_2_-induced p53 activation. RPE^Tert^ cells were pre-treated with DMSO or PH797804 (10 μM) for 2h, followed by treatment with diamide (200 μM) or H_2_O_2_ (200 μM) for the indicated time. The total protein levels of p38MAPK, p53, p21 and GAPDH, and phosphorylation states of p38MAPK(T180/Y182), ATF-2(T71) and p53(S33) were evaluated by immunoblotting. (D) RPE^Tert^ cells were pre-treated with DMSO or SP600125 (20 μM) and PH797804 (10 μM) together for 2h, followed by treatment with diamide (200 μM) or H_2_O_2_ (200 μM) for the indicated time. The total protein levels of JNK (p54 and p46), p38MAPK, p53, p21 and GAPDH, and phosphorylation states of JNK(T183/Y185) and p38MAPK(T180/Y182), were evaluated by immunoblotting analysis.

### 3.5 Redox signaling activates p53-dependent transcriptional activity

The above presented data (see e.g. Fig. 2 and 4) already show that activation of redox signaling either by diamide or H_2_O_2_ treatment induces p21 expression in a p53 dependent manner. The notion that DDR and redox signaling dependent p53 activation proceeds through different upstream kinase signaling cascades, could in principle lead to an induction of different p53 transcriptional targets due to alternative PTM or cofactor binding. In order to evaluate p53-dependent gene transcription we established doxycycline-inducible Flag-p53 expressing p53 KO RPE^Tert^ cells. Doxycycline (dox) treatment was optimized to induce Flag-p53 levels similar to endogenous p53 in the parental RPE^Tert^ cell line under basal conditions (4 ng/ml dox treatment for 48 h or 72 h, **Fig. 5A**). Ectopically expressed Flag-p53 in these cells mimicked the response to diamide and H_2_O_2_ observed in wildtype RPE^Tert^ cells (**Fig. 5B**). Next, we evaluated the expression of *TP53* itself and some target genes of p53 associated with cell cycle arrest (*CDKN1A* and *GADD45a*), apoptosis (*BAX* and *PIG3*), p53 turnover (*MDM2*) and metabolism (*TIGAR*) upon redox signaling and DNA damage. We found that *TP53* mRNA levels were increased by almost 2-fold upon addition of doxycycline, and was significantly induced further by H_2_O_2_ treatment, suggesting that H_2_O_2_ regulates p53 levels both at the level of transcription and stabilization **(Fig. 5C and Fig. 3A&B)**. p53 transcriptional targets, *CDKN1A* (p21), *GADD45a* and *PIG3* were further activated both by redox signaling and DDR signaling to p53 to some extent, whereas *MDM2 and BAX* were significantly induced only by DNA damage signaling to p53 (H_2_O_2_ and NCS). No obvious additional change in the induction of *TIGAR* was observed upon either treatment. Collectively, these observations suggest that both redox signaling and the DDR can activate p53, and that there seems to be some target selectivity depending on which upstream pathway activates p53.

**Figure 5.**
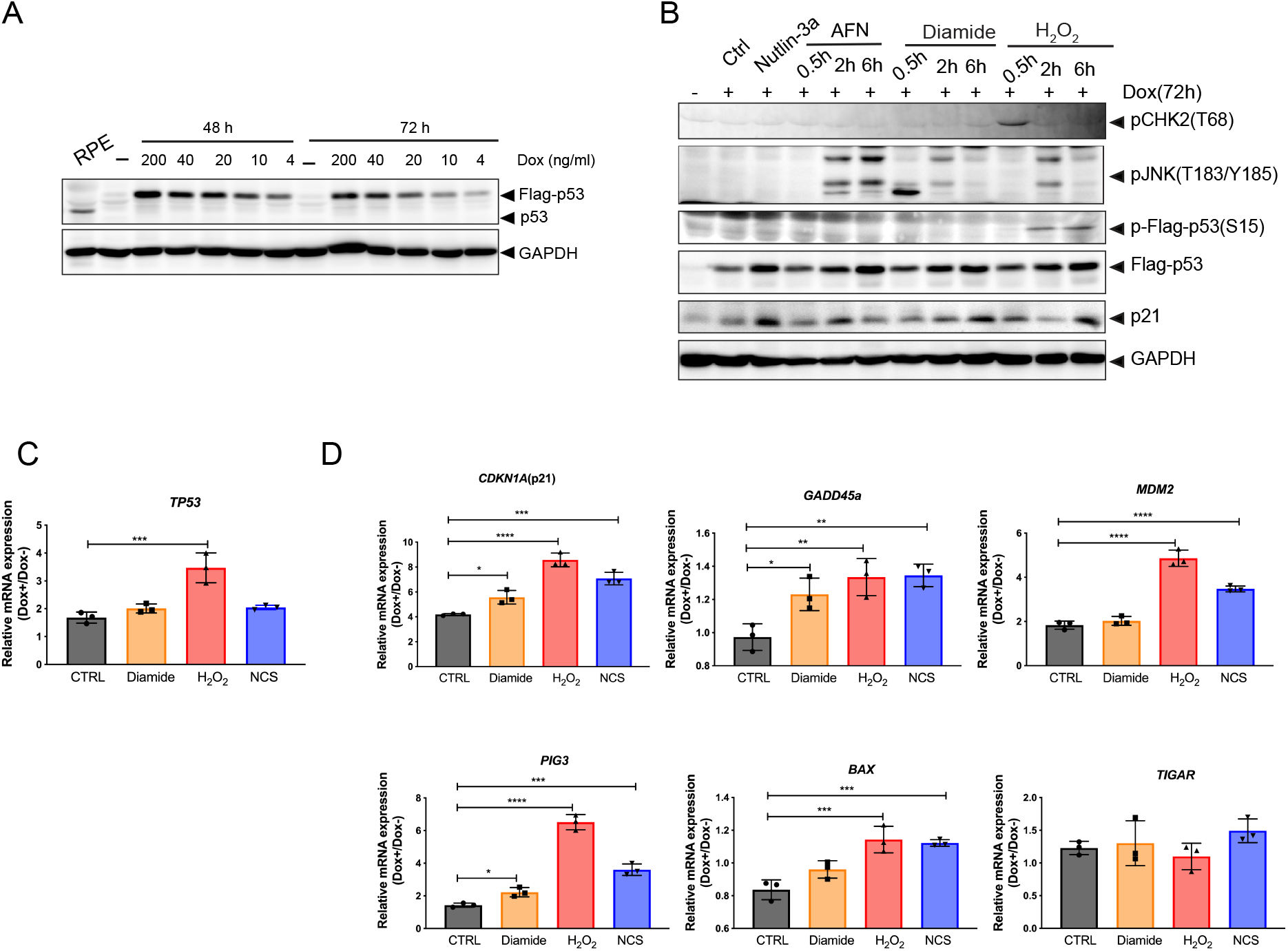
Redox signaling induces p53-dependent transcriptional activation. (A) Immunoblotting analysis of RPE^Tert^ p53 KO cells expressing Dox-inducible p53, treated with a range of doxycycline (dox, 4-200 ng/ml). Wildtype p53 in RPE^Tert^ cells is used as a reference for endogenous levels. (B) p53 expression was induced with 4 ng/ml Dox for 72h to mimic near-endogenous levels, followed by treatment with Nutlin-3a (10 μM), Auranofin (AFN) (10 μM), diamide (200 μM) or H_2_O_2_ (200 μM) for the indicated time. Total cell lysates were analyzed for the levels of Flag-p53, p21 and GAPDH, and phosphorylation states of CHK2(T68), JNK(T183/Y182) and Flag-p53(S15) by immunoblotting. (C) p53 expression was induced with 4 ng/ml Dox for 48h, followed by treatment with diamide (200 μM), H_2_O_2_ (200 μM) or NCS (500 ng/ml) for 24 h. The expression of p53 target genes was measured in both Dox− (p53-off) and Dox+ (p53-on) cells by qPCR. The ratio of the gene expression (relative to GAPDH) in p53-on cells over that in p53-off cells was calculated to assess p53-dependent transcriptional target activation. The data is presented as the mean and standard deviation (SD) from three independent experiments. Statistical analysis was performed by using One-way ANOVA followed by Multiple comparisons test (Dunnett). *, p value < 0.05; **, p value < 0.005; ***, p value < 0.0005; **** p value < 0.0001.

### 3.6 Both Redox signaling and DNA damage trigger p53-dependent cell death

To further examine the biological consequences of p53 activation by redox signaling and DNA damage, we examined p53-dependent cell death in RPE^Tert^ p53 KO cells expressing doxycycline inducible p53 upon treatment with diamide, H_2_O_2_ or NCS for 24 hrs (**Fig. 6A&B**). We observed that both diamide and H_2_O_2_ treatment induced significantly more cell death following p53 expression, indicating that both the DDR and redox signaling can trigger p53-dependent cell death. Note that NCS did not induce cell death in this cell line, whereas it did in H1299 cells **(Supple. Fig. 1A&B)**, which could be in line with the general notion that cancer cells are more vulnerable to chemotherapeutic drugs than untransformed cells **[32]**. Both diamide and H_2_O_2_ also induced p53-dependent cell death in H1299 cells **(Supple. Fig. 1A&B)**. Collectively, our data reveal that redox signaling can activate p53 to induce cell death in the absence of the DDR both in untransformed and human cancer cells.

**Figure 6.**
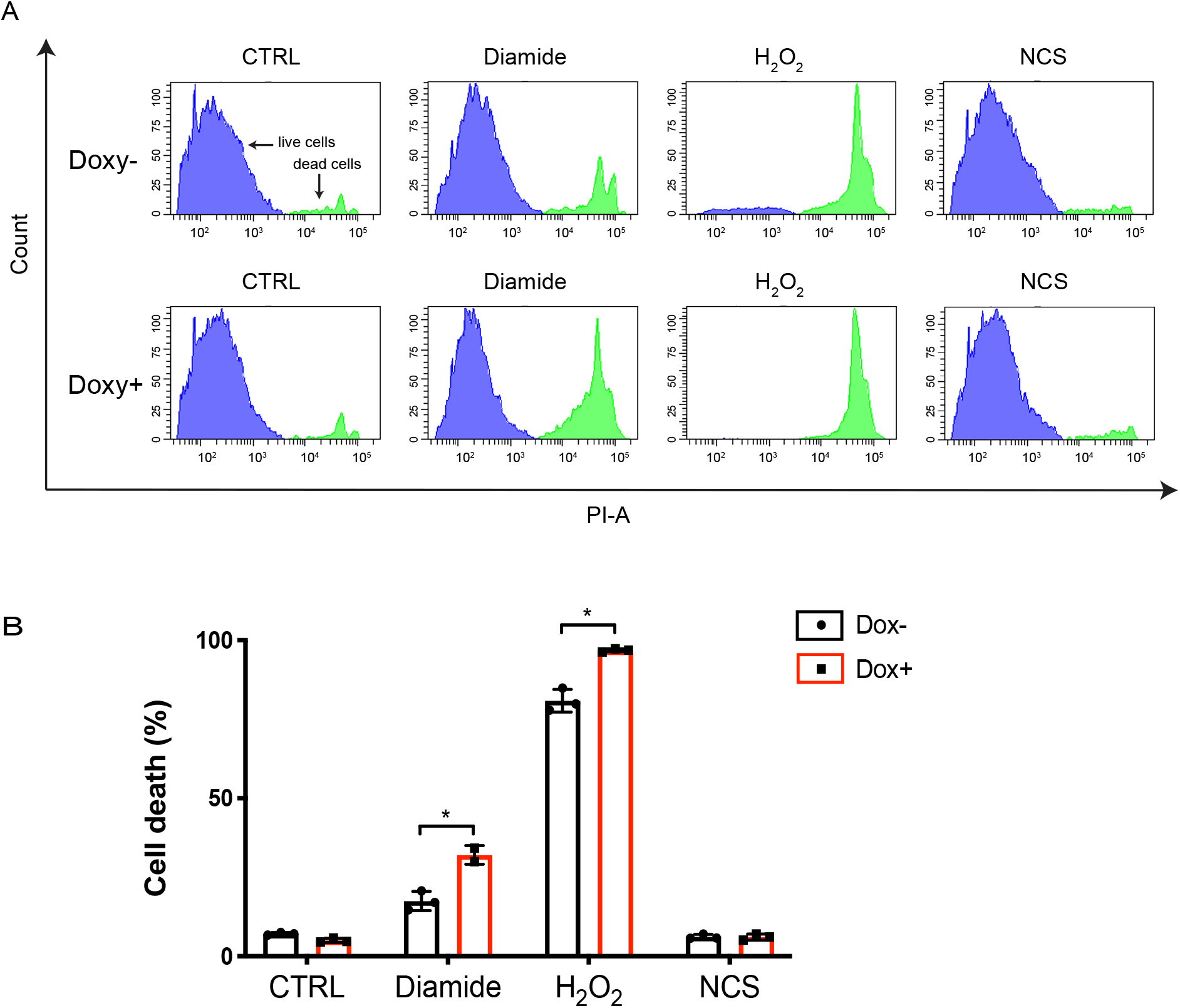
Redox signaling and DNA damage induce p53-dependent cell death. (A) Histogram plots showing cell death in Dox inducible expressing p53 RPE^Tert^ cells upon induction of redox and DNA damage signaling as measured by Flow Cytometry (PI-exclusion assay). Dox-inducible expressing p53 RPE^Tert^ cells were cultured with or without Dox for 48 h, followed by the addition of diamide (250 μM), H_2_O_2_ (300 μM) and NCS (500 ng/ml) for 24 h. Cell death was then measured by Flow Cytometry using Propidium iodide (PI) staining. The plots show representative samples from three independent experiments. (B) Quantification of cell death from three independent experiments. A student’s t-test was used to analyse statistical difference in cell death between Dox− and Dox+ RPE^Tert^ cells upon each treatment. *, p value < 0.05.

## 4. Discussion

The observations that exposure to ROS (H_2_O_2_, O_2_^•−^, HO^•^) either from endogenous or exogenous sources can activate the DDR as well as p53 **[4, 13, 33]** has given credence to the idea that ROS activates p53 downstream of signaling in response to oxidative DNA damage. In line with this notion, (enhanced) mitochondrial respiration and the ensuing O_2_^•−^/ H_2_O_2_ production is frequently cited as a source of oxidative DNA damage and mutation in genomic DNA in tumors **[16, 17]**. But H_2_O_2_ also acts as a second messenger in redox signaling, which plays an essential role in regulating protein functions and biological processes **[18]**, including several phosphorylation cascades upstream of p53 **[19, 20]**. As outlined in the introduction section, it has therefore not been straightforward to attribute p53 activation in response to elevated H_2_O_2_ levels to activation of the DDR, redox signaling or both **[10]**. What further complexes understanding ROS-induced p53 activation is the observation that ATM can also be activated by redox signaling in the absence of DNA damage **[34]**. Furthermore, treatment with several DNA damaging chemotherapeutic drugs, such as doxorubicin, cisplatin and 5-fluorouracil, can lead to enhanced ROS production **[35, 36]**. In this work, we set out to dissect DNA damage signaling and redox signaling upstream of p53, by applying treatments that we carefully titrated and validated to either induce only the DDR (as judged by gamma-H2AX and pCHK2), only redox signaling (as judged by oxidation of the H_2_O_2_ specific HyPer2) probe or both. NCS induced a DNA damage response, without evidence of elevated H_2_O_2_ within 6 h. Diamide only led to redox signaling without activation of the DDR, whereas H_2_O_2_ indeed induced both DDR and redox signaling, each with similar kinetics as observed for NCS and diamide respectively.

Other studies did find that treatment with NCS resulted in the elevated oxidation of 2,7-dichlorodihydrofluorescein (DCFH_2_-DA) in U2OS cells **[37]**. The different cell lines used, specificity of the used detection method or drug concentrations applied may underlie this apparent discrepancy. Besides the lack of HyPer oxidation induced by NCS, we also found no evidence of NCS-induced redox signaling as judged by p38MAPK or JNK activation in RPE, H1299 and HEK293T cells (not shown) at the concentration range used in this study, which we think further validates the approach. As mentioned, it has been shown that ATM can be activated in the absence of DNA damage through disulfide-dependent homodimerization in response to oxidant treatment **[21]**. In contrast to our findings, that study found that both H_2_O_2_ and diamide were capable of disulfide-dependent activating ATM and subsequent p53-Ser15 phosphorylation, whereas we found no evidence that diamide could activate ATM as judged by p53-pSer15, CHK2-Thr68 or gamma-H2AX in all cell lines we tested. On the other hand, the same study found induction of pCHK2 but not gamma-H2AX in response to H_2_O_2_, whereas in our study H_2_O_2_ did induce both pCHK2 and gamma-H2AX as well as activation of p38MAPK and JNK, suggesting that H_2_O_2_ triggers both the canonical DDR along with redox signaling. Again, differences in the precise protocol for treatment and cell lines used might underlie these contrasting observations.

With the described careful selection of treatments that trigger only the DDR, only redox signaling or both we were able to dissect how p53 is activated upstream by these pathways. We observed that p53 was activated by DNA damage and redox signaling through distinct upstream kinases, that both seem to converge on the inhibition of MDM2-dependent p53 degradation **(Fig.7)**. ATM was required for p53 activation in response to NCS and H_2_O_2_-induced DNA damage, but dispensable for p53 activation induced by diamide-mediated redox signaling. p38MAPK activity on the other hand was required for p53 stabilization and activation upon diamide-induced redox signaling, but not for NCS and H_2_O_2_-induced p53 activation. Our results furthermore showed that JNK was required for p53 activation by H_2_O_2_, although inhibition of ATM also completely blocked H_2_O_2_-induced p53 activation. This could suggest that ATM activation also somehow requires JNK activity in case of H_2_O_2_ dependent activation, although this remains to be further explored. H_2_O_2_ activates both JNK and p38MAPK, which suggests that H_2_O_2_ treatment would still result in p53 stabilization in the presence of JNK inhibitor through the p38MAPK pathway, but we did not clear find evidence for this. This could be because the extent of p38MAPK activation by H_2_O_2_ is much lower as compared to diamide dependent activation, or there might be other undiscovered pathways induced by diamide but not H_2_O_2_ that act in concert with p38MAPK.

**Figure 7.**
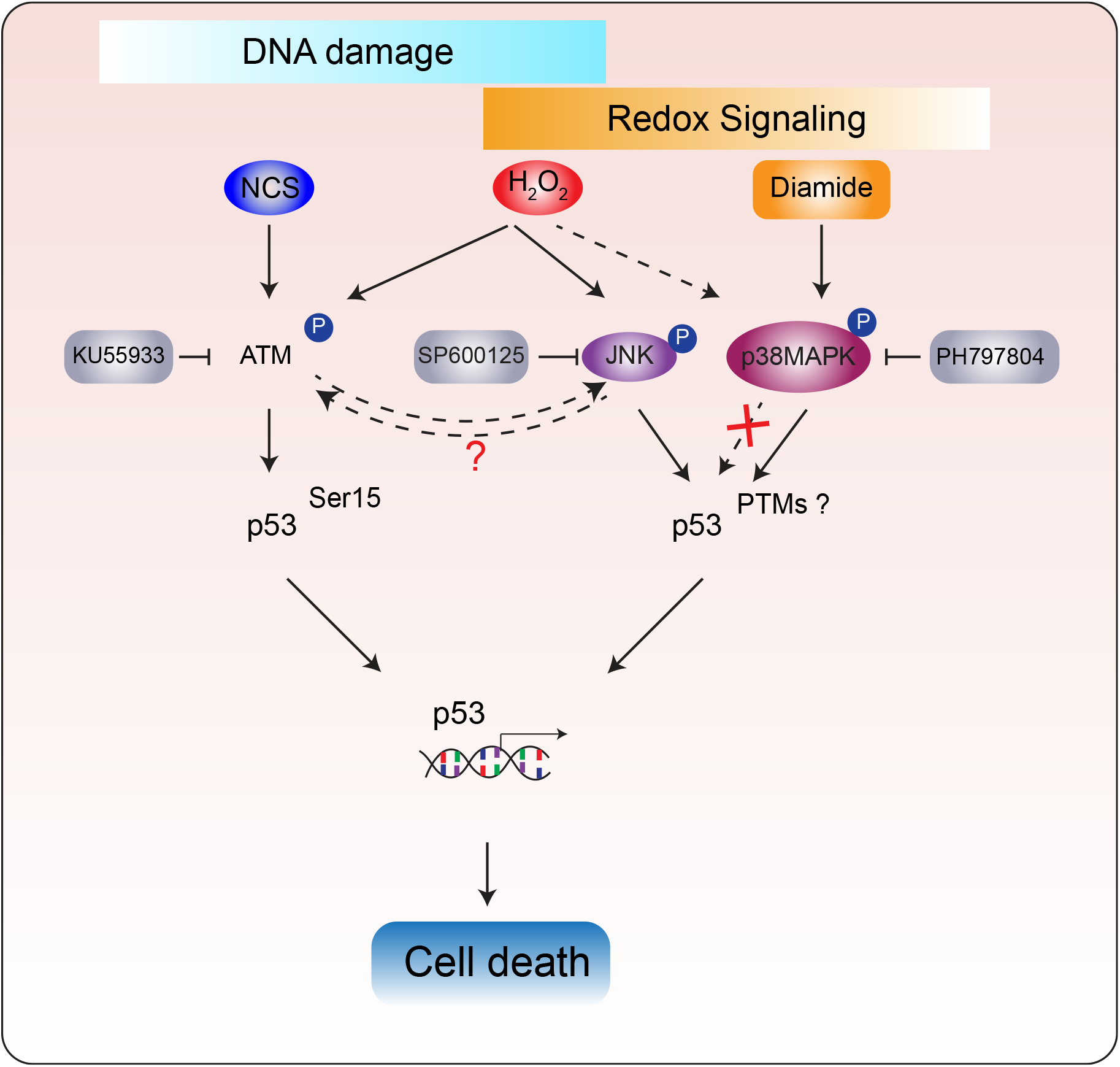
Distinct upstream kinase-dependent signaling pathways activate p53 in response to DNA damage and redox signaling. NCS and H_2_O_2_ both trigger the ATM-dependent DNA damage response and downstream p53 activation. Inhibition of ATM indeed abolishes NCS and H2O2-induced p53 activation. H2O2 also activates JNK, and this was also required for H_2_O_2_-induced p53 activation, whereas inhibition of JNK abrogated p53 activation by H2O2, no effects on the ATM pathway were observed, suggesting that JNK and ATM somehow mediate p53 activation through synergystic pathways, but the details remains to be unraveled. p38MAPK is also activated by H2O2, but it was required for H2O2-induced p53 activation (dashed line). In contrast, diamide-induced redox signaling activates p53 through p38MAPK, independent of ATM and JNK. p53 induces transcriptional target genes and cell death in response to both the DDR and redox signaling. Our data indicate that p53 can be activated by redox signaling without inducing collateral DNA damage, thereby lowering the risk for the acquisition of new mutations driving tumor progression and initiation

We found that activation of p53 by both redox signaling and the DDR resulted in transcriptional activation of p53 targets, and there seemed to be some differential effects dependent on which pathways were activated. It has been proposed that different stresses would lead to distinct transcriptional programs of p53, including in response to redox signaling and DNA damage **[38]**. Differential regulation in response to specific stressors could stem from alternative co-factor binding, specific PTMs or the simultaneous engagement of parallel signaling pathways, and it has also been suggested that oxidant-induced p53 target gene promoters bear distinct p53 consensus motifs **[39].** However, in our study we did not observe a black and white of p53-dependent gene expression in response to differential stresses. We found that p21, *GADD45a* and *PIG3* induced by both redox signaling and DNA damage, whereas *MDM2* and *BAX* were more induced by DNA damage signaling to p53 (H_2_O_2_ and NCS) than redox signaling (diamide). Since most previous studies did not carefully compare the induction of target genes in response to compounds that only induce the DDR or redox signaling, it is difficult to compare our observations to these studies. Furthermore, our selection of p53 target genes is rather limited and mostly aimed at showing that both DDR and redox signaling induced p53 stabilization also activates its transcriptional activity.

Arguably the prime p53-dependent tumor suppressive response is the induction of apoptosis, which is the goal of many anti-cancer therapies that are aimed at the reactivation or restoration of wild-type p53 function. Primary cancer-therapies including several chemotherapeutics and irradiation elicit DNA damage and trigger the DDR and downstream p53-dependent apoptosis in multiple tumor types **[40, 41]**. Some chemotherapeutics used in the clinic, like Cisplatin and Doxorubicin have been shown to activate both JNK and p38MAPK along with the DDR**[7, 42]**, but it is not entirely clear which of these pathways represents the dominant mechanism behind their efficacy. But the induction of DNA damage comes with the risk of generating new mutations. These may induce novel oncogenic events in surrounding tissue, but also drive tumor progression and therapy resistance through tumor evolution by mutation and selection. The data presented here suggest that p53 can be activated to trigger an apoptotic response independent of the DDR through redox signaling. This implies that it would in theory be possible to develop therapeutics that restore a p53 tumor suppressive response without risking the induction of collateral DNA damage and the ensuing tumor cell evolution.

Tumor cells, as compared to healthy cells, in general have higher ROS levels through, for instance altered metabolism, and as a result need to augment their antioxidant capacity in order to survive and thrive. It has been suggested that due to the simultaneous elevated production and scavenging of ROS in tumor cells, the redox state would be more easily tilted to more oxidizing **[43]**. With that in mind, further enhancing ROS levels using pro-oxidant approaches have been suggested as a strategy to induce tumor cell death **[44]**. But inhibition of the cellular reductive capacity, like we do here using diamide, could in principle trigger a p53 response without the risk of collateral DNA damage as explained above. Such therapies may for instance be aimed at inhibition of the TrxR/Trx system (using e.g. Auranofin) **[45, 46]** or depletion of NADPH **[47]** but it remains to be explored whether such approaches would indeed be feasible. Taken together, the observation that redox signaling and the DDR activate p53-apoptosis through distinct upstream signaling cascades may contribute to new ideas for developing therapeutic strategies.

## Author contributions

T.S. and T.B.D. designed the study and wrote the manuscript. B.M.T.B. reviewed and commented on the manuscript. T.S. performed most of the experiments. P.E.P established the doxycycline-inducible Flag-p53 expressing system in cells and assisted with laboratory management.

## Acknowledgments

We are grateful for suggestions and input from our colleagues at the department of Molecular Cancer Research, University Medical Center Utrecht. We thank Jeroen van den Berg and René Medema for sharing their RPE^Tert^ p53 KO cell line, Vsevolod Belousov for sharing the HyPer2 construct and Daan van Soest for help with using this probe. The work was made possible with grants from the China Scholarship Council (CSC no. 201606300046) to T.S. and from the Dutch Cancer Society (KWF UU 2014-6902) to T.B.D. B.M.T.B is part of the Oncode Institute, which is partly financed by the Dutch Cancer Society (KWF Kankerbestrijding) and was funded by the gravitation program CancerGenomiCs.nl from the Netherlands Organization for Scientific Research (NWO).

---

**Supplemental Figure S1. Redox signaling and DDR induced, p53-dependent cell death in H1299 cells**

**(A)** Dox-inducible p53 expressing H1299 cells were cultured with or without Dox for 48 h, followed by treatment with diamide (200 μM), H_2_O_2_ (500 μM) and NCS (500 ng/ml) for 24 h. Cell death was then measured by Flow Cytometry using Propidium iodide (PI) staining. The data is from a representative sample from two independent experiments.

(C) Quantification of cell death from two independent experiments. A student’s t-test was used to analyse statistical difference of cell death between Dox− and Dox+ H1299 cells upon each treatment. p value < 0.05 is considered to be statistically significant. Bars show mean and SD of two independent experiments.

