## Supplemental Figure S1 for "DNA damage and oxidizing conditions activate p53 through differential upstream signaling pathways"

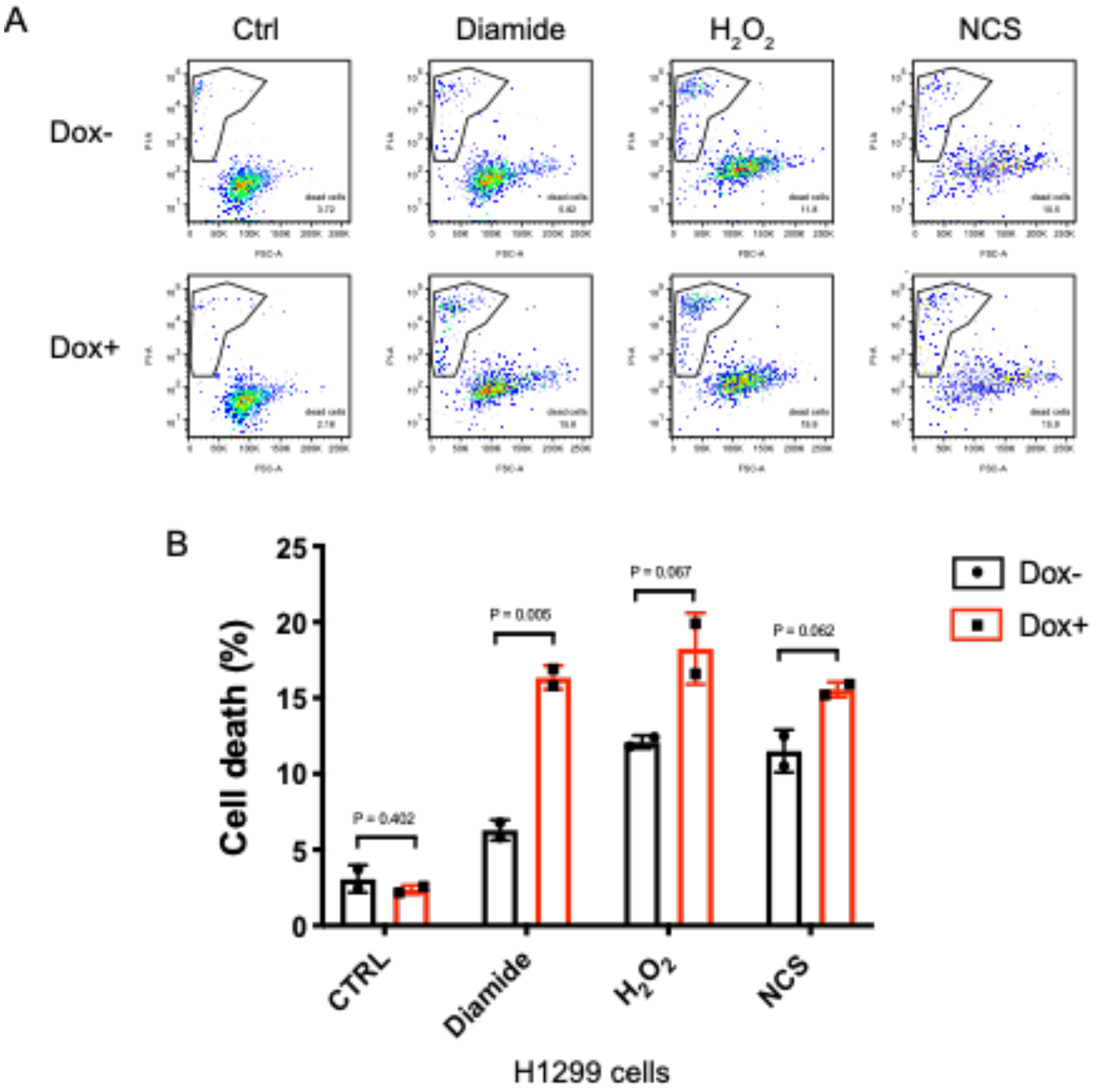


**Supplemental Figure S1. Redox signaling and DDR induced, p53-dependent cell death in H1299 cells**

(A) Dox-inducible p53 expressing H1299 cells were cultured with or without Dox for 48 h, followed by treatment with diamide (200 µM), $H_2O_2$ (500 µM) and NCS (500 ng/ml) for 24 h. Cell death was then measured by Flow Cytometry using Propidium iodide (PI) staining. The data is from a representative sample from two independent experiments.

(B) Quantification of cell death from two independent experiments. Bars show mean and SD of two independent experiments. A student's t-test was used to analyse statistical difference of cell death between Dox- and Dox+ H1299 cells upon each treatment. p value < 0.05 is considered to be statistically significant.
