## Supplemental Table S1 for "DNA damage and oxidizing conditions activate p53 through differential upstream signaling pathways"

| Gene | sequences(5'-3') | References |
|---|---|---|
| GAPDH-For | CATTTCCTGGTATGACAACG | |
| GAPDH-Rev | CTCTTCCTCTTGTGCTCTTG | |
| TP53-For | CTCAGATAGCGATGGTCTGG | |
| TP53-Rev | CAAATACTCCACACGCAAAT | |
| CDKN1A-For | AGCAGGCTGAAGGGTCCCCA | |
| CDKN1A-Rev | GGCGTTTGGAGTGGTAGAAATCTGT | |
| GADD45a-For | GAGAGCAGAAGACCGAAAGGA | zhang et al., 2011 |
| GADD45a-Rev | CAGTGATCGTGCGCTGACT | |
| MDM2-For | CAGGCAAATGTGCAATACC | |
| MDM2-Rev | GTGCACCAACAGACTTTAAT | |
| PIG3-For | CCGGAAAACCTCTACGTGAA | Kung et al., 2014 |
| PIG3-Rev | ATGCCTCAAGTCCCAAAATG | |
| BAX-For | CCCGAGAGGTCTTTTTCCGAG | |
| BAX-Rev | CCAGCCCATGATGGTTCTGAT | |
